# Amino acid and carbohydrate tradeoffs by honey bee nectar foragers and their implications for plant-pollinator interactions

**DOI:** 10.1101/008516

**Authors:** Harmen P. Hendriksma, Karmi L. Oxman, Sharoni Shafir

**Affiliations:** B. Triwaks Bee Research Center, Department of Entomology, Robert H. Smith Faculty of Agriculture, Food and Environment. The Hebrew University of Jerusalem, Rehovot 76100, Israel

**Keywords:** pollination ecology, homing, essential amino acids, equivalence point, nutrient balance, pH deviation

## Abstract

Honey bees are important pollinators, requiring floral pollen and nectar for nutrition. Nectar is rich in sugars, but contains additional nutrients, including amino acids (AAs). We tested the preferences of free-flying foragers between 20 AAs at 0.1% w/w in sucrose solutions in an artificial meadow. We found consistent preferences amongst AAs, with essential AAs preferred over nonessential AAs. The preference of foragers correlated negatively with AA induced deviations in pH values, as compared to the control. Next, we quantified tradeoffs between attractive and deterrent AAs at the expense of carbohydrates in nectar. Bees were attracted by phenylalanine, willing to give up 84 units sucrose for 1 unit AA. They were deterred by glycine, and adding 100 or more units of sucrose could resolve to offset 1 unit AA. In addition, we tested physiological effects of AA nutrition on forager homing performance. In a no-choice context, caged bees showed indifference to 0.1% proline, leucine, glycine or phenylanaline in sucrose solutions. Furthermore, flight tests gave no indication that AA nutrition affected flight capacity directly. In contrast, low carbohydrate nutrition reduced the performance of bees, with important methodological implications for homing studies that evaluate the effect of substances that may affect imbibition of sugar solution. In conclusion, low AA concentrations in nectar relative to pollen suggest a limited role in bee nutrition. Most of the 20 AAs evoked a neutral to a mild deterrent response in bees, thus it seems unlikely that bees respond to AAs in nectar as a cue to assess nutritional quality. Nonetheless, free choice behavior of foraging bees is influenced, for instance by phenylalanine and glycine. Thus, AAs in nectar may affect plant-pollinator interactions and thereby exhibit a selective pressure on the flora in the honey bee habitat.

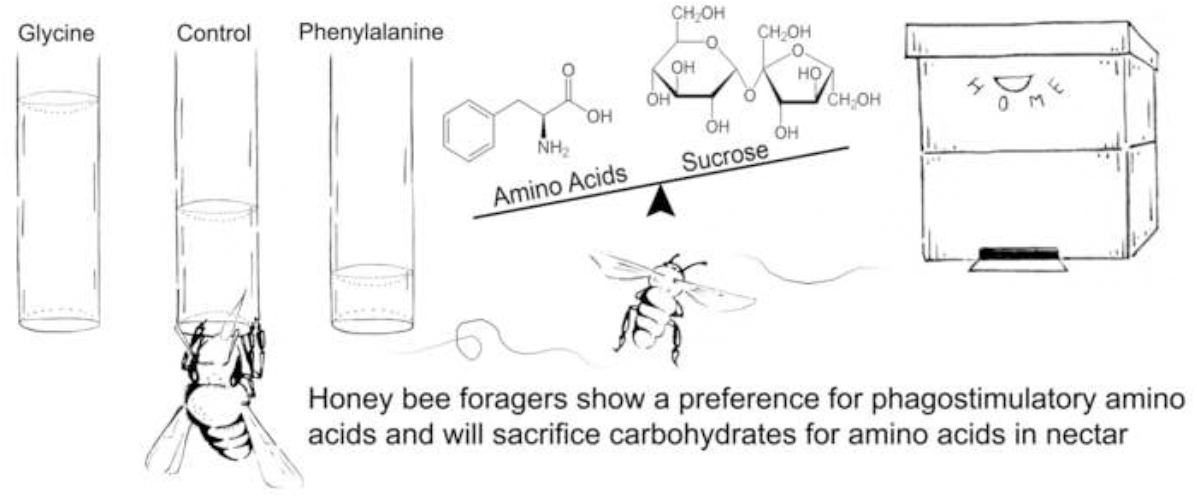

**HIGHLIGHTS:**

– Amino acids in artificial nectar elicit preferences from honey bee foragers
– Amino acid identity, pH, and essentiality explain preferences of bees
– A honey bee forager is willing to pay a premium of carbohydrates for amino acids
– Carbohydrate nutritional state affects flight performance of foraging bees

## 1. Introduction

The honey bee, *Apis mellifera,* is a key contributor to the pollination of crops worldwide. The nutritional need of honey bee colonies for floral pollen and nectar is the fundamental driver for this valuable ecosystem service. Pollen contains nutrients for growth, such as essential amino acids (AAs) in proteins, certain lipids, and essential trace elements like minerals and vitamins (Free, 1987; Herbert and Shimanuki, 1978). Nectar is the main source of sugars and additionally it provides AAs, lipids, antioxidants, and potentially toxic secondary metabolites (Baker, 1977). The proportion of forager bees for pollen and nectar is determined by the nutritional state of the colony and the availability of floral resources in the environment (Calderone and Johnson, 2002; Camazine, 1993; Seeley, 2009).

It is still debated whether pollen-collecting insects, such as honey bees (Fewell and Winston, 1992; Saa-Otero et al., 2000) and bumblebees (Rasheed and Harder, 1997a, b), maximize foraging efficiency by preferentially switching to protein-rich pollen. Whereas Levin and Bohart (1955) found for five of six pollen types offered at feeding stations that the preference ranked according to crude protein content, Schmidt (1982) found no clear relation between preferences for eight pollens and their protein content. Similarly, choice experiments have shown that honey bees do not assess pollen protein content (Pernal and Currie, 2001, 2002). This is not surprising considering that protein is very rarely present in the pollenkitt, the oily layer coating the pollen grain, and free amino acids were not found in the pollenkitt of any of 69 plants studied (Dobson, 1988). It appears that cues other than protein content affect bee pollen preferences. Such cues may include odors, phagostimulants and phagodeterrents (Schmidt, 1982), grain size (Pernal and Currie, 2002), and pollen concentration; honey bees dance more rigorously to a feeder containing a higher pollen-to-cellulose ratio (Waddington, 2001; Waddington et al., 1998).

In contrast, far more is known on the ability of forager bees to evaluate the nutritional value of nectar. The main total dissolved solids (TDS) in nectar are sugars, and bees can readily determine sugar content. They can discriminate small differences in sugar concentration (Afik et al., 2006; Frisch, 1967; Shafir et al., 2008), and nectar volume and variability (Shafir et al., 2005; Waddington and Gottlieb, 1990). Honey bees are able to perceive and show preference for nectar mineral contents, secondary compounds and certain AAs (Afik et al., 2006; Carter et al., 2006; Cook et al., 2003; Gardener and Gillman, 2002; Kim and Smith, 2000; Singaravelan et al., 2005). The means of sensing nectar composition are receptor based, with different kinds of taste receptors on the proboscis and mouth parts (Goodman, 2003; Sanchez, 2011). It is unknown however, to what extent amino acid (AA) presence in nectar is valued, and further, if potential preferences are related to the nutritional state of the colony. Whereas pollen is the main source of AAs for bees, we cannot exclude the possibility that deficiencies in particular AAs in the colony may also modulate preference for nectars which contain those AAs.

A colony needs a minimum level of essential AAs to assimilate proteins for growth (De Groot, 1953). Hence, a preference for essential over non-essential AAs may be an adaptive strategy for bees. To date however, a general preference for essential AAs has not been found, though certain individual AAs are known to evoke phagostimulant or phagodeterrant effects on bees (Inouye and Waller, 1984). Honey bee studies have tested AAs singly, e.g. on consumptive responses (Inouye and Waller, 1984) or on olfactory perception (Linander et al., 2012). However, in the present study, we offered all AAs simultaneously in order to compare AA preferences in relation to one another. This may reveal insights in how honey bees in a floral habitat respond to a complexity of choice options. An AA-induced gustatory preference may influence plant pollinator interactions, which can have ecological and evolutionary consequences by affecting gene flow within and between plant populations (Gardener and Gillman, 2002; Gottsberger et al., 1984; Nepi et al., 2012). In addition to direct preference effects, AAs may induce more subtle plant pollinator interactions. AA levels in bees may stimulate or inhibit learning and memory (Chalisova et al., 2011), thereby influencing flower association by free flying foragers, as has been shown for nicotine in nectar (Wright et al., 2013).

Forager preferences for AAs may come with benefits. Proline for example, has been described as a flight muscle stimulant (Barker and Lehner, 1972; Carter et al., 2006; Micheu et al., 2000; Mollaei et al., 2013). At the same time, a preference may come at a cost when the perceived profitability exceeds the benefit. This would be the case if uptake due to phagostimulating AAs were in lieu of carbohydrate uptake.

If foragers alter their intake of carbohydrates, they might pay a physiological price, either in terms of fuel for flight, or their flight ability could change, for instance due to alterations in their carrying load. By means of homing experiments, we explored whether AAs in nectar elicit physiological effects on forager bees. In particular, we compared the indirect effect of phagostimulant and phagodeterent AAs on sucrose uptake, consequent nutritional state of the bee, and its effect on homing success and flight speed.

An additional aim of our study was to test the relative preferences of free flying nectar foragers between simultaneously presented AA-enriched sucrose feeders. Since AAs in artificial nectar may affect forager choice, we tested for a tradeoff between AAs and carbohydrate in collected nectar. We quantified evoked responses by comparing units of AAs to equivalent units of sugar, thereby calculating the energetic cost of choice.

## 2. Materials and Methods

The experiments were conducted at the Benjamin Triwaks Bee Research Center in Rehovot, Israel, with the local line based mostly on the Italian honey bee strain *Apis mellifera ligustica*.

### 2.1. Preference between nonessential and essential amino acids

In March, October and November 2013, bees were trained to forage on ten 200 ml feeders filled with honey-tainted sucrose solutions (30% w/w), hanging at fences in the vicinity of two of our apiaries. At any time, 20 to 30 colonies of free-flying bees were present. Once foragers were visiting the fence sites, a few dozen 2 ml glass tubes with one open end were filled with 20% sucrose solutions and hung on 2.5 x 2 m fence sections to train the bees to visit the tubes. The feeding tubes together on the fence formed an artificial meadow to conduct experiments. In order to unravel the complexity of nectar foragers choice patterns, this novel test approach allows testing a wide range of treatments.

To test honey bee forager preferences among AA-enriched artificial nectars, a series of choice experiments was started by hanging new tubes with AA solutions in the fence sections. Each AA treatment was presented in 1.8 ml 20% sucrose solutions, enriched with an AA at 0.1% (w/w). Abbreviations for amino acid names used throughout the manuscript are listed in Supplement 2. Depending on molecular weights of each AA, treatment solutions were in range of 5.3 – 14.3 mM (Supplement 2). The solutions were always offered in presence of a control (20% sucrose solution). Considering a crop volume of maximally 70 µL, it takes at least 25 bee visits to empty a tube with a volume of 1.8 ml of liquid.

In three experimental designs, we tested 8 nonessential AAs (72 test tubes offered to bees in 8 replicate fence sections), 10 essential AAs (88 test tubes in 8 replicate sections), and 20 AAs together (542 feeding tubes in 25 replicate sections). The tubes were hung according a Latin square design, so that each treatment appeared in a different spatial position in every replicate fence section. The experiment with all the AAs was replicated at March 13 and 15, and November 16, at two apiaries. Each apiary contained 20-30 hives, which were exchanged from time to time. Additional details of the experimental designs are given in the supplementary material (Supplement 1: S1-a, S1-b and S1-c). The pH values of the solutions were measured for each batch of test solutions (Supplement 2).

The fluid levels in the test tubes were marked at the meniscus at the start and end of the experiment. As soon as the first tubes in a section were nearly or completely empty, all tubes in that section were turned upwards, so that the content was no longer available to the bees. We then measured the column height of the collected solution in each glass tube to the nearest mm. For every AA treatment, proportional consumption values were calculated on the total solution collected per replicate fence section.

### 2.2. Tradeoff between carbohydrates and amino acids

To quantify the tradeoff between sucrose and amino acids, glass tubes were hung at 12 replicate fence sections, offering 12 AA treatments to free flying foragers. The test was conducted as a 12x12 Latin square design (Supplement 1: S1-d). Treatments were control solutions of 15, 18, 21, and 24% sucrose (w/w), solutions of 15, 18, 21, and 24% sucrose with 0.1% phenylalanine (Phe), and solutions of 15, 18, 21, and 24% sucrose with 0.1% glycine (Gly). Based on the findings from the AA-choice experiment (see 2.1.), phenylalanine was chosen to represent a phagostimulant, and glycine to represent a phagodeterrent. The phenylalanine and glycine treatments had pH values similar to the control (pH_Control_ = 6.15, pH_Phe_ = 6.23, pH_Gly_ = 6.28). Data were collected on the proportional consumption of treatment solutions per fence section. The data were then used to extrapolate sucrose tradeoff levels where a 0.1% AA solution would be equally collected compared with a sucrose control.

We tested these extrapolated values in two additional experiments. In one experiment, there were three treatments: a 20% sucrose control, a lower sucrose concentration with 0.1% phenylalanine, and a higher sucrose concentration with 0.1% glycine. Throughout 9 fence sections, in which each solution was represented twice, 18 replicates of each solution were presented to free-flying bees (Supplement S4). The results confirmed the equivalence sucrose concentration for phenylalanine, but suggested a higher sucrose equivalence concentration for glycine. In a follow-up experiment, we presented four treatments: a 20% sucrose control, and 20%, 30% and 40% sucrose solutions each enriched with 0.1% glycine (Supplement S4).

### 2.3. The effect of amino acid and carbohydrate consumption on homing success

To study AA and sucrose nutrition effects on foragers, three homing tests were performed (H1, H2 and H3). All test individuals were homecoming foragers that were collected by closing the entrance to a test hive. Captured bees were placed on ice for several minutes until they stopped moving, marked on their thorax with a color to indicate the treatment they would receive, and placed in groups of 15 or 20 bees in a clear plastic jar with a mesh opening on the side for ventilation. Over the three homing experiments, a total of five hives were tested, with a total of 2079 bees placed in 130 clear plastic jars (details in Supplement 3).

Every jar received two glass tubes through holes in the lid, each containing 1.8 ml of test solution. The jars were then placed in an empty hivebox, on top of a populated hivebox. The two boxes were separated by means of a queen excluder covered with a 1 mm2 mesh. This allowed the bees in the jars above to feel the warmth and scent of the colony below. The levels of test solution were marked on the tubes as soon as they were inserted into the jars. The levels were marked twice a day, in parallel to noting the number of dead bees per jar, to determine how many individuals contributed to consuming the treatment solutions over time, and to monitor for treatment-related mortality effects.

On the day of the homing experiment, bees were released forenoon approximately 600 m from the hive. Bees were released onto an arena. During the thirty minutes after release, moribund and dead individuals were collected and counted. The “dead” bees included those that died during the pre-phase of the experiment, whereas “moribund” bees were those that were not able to fly away from the arena within thirty minutes. Experimenters back at the test hives were notified when the bees were released. The hive entrances were closed and every homecoming marked bee was captured, in order to prevent recounts, over a three hour time period. The return time and color for every test bee was recorded. After each experiment, the test hives were opened to verify the absence of marked bees.

During the first homing experiment (H1), the treatments were 40% sucrose (1), and 40% sucrose enriched by 0.1% phenylalanine (2), 0.1% leucine (3), 0.1% proline (4), or 0.1% glycine (5). Alike the fence experiments, these particular amino acids were chosen to represent phagostimulants, phagodeterrents, nonessential, and essential amino acids as well as being pH neutral in comparison to the control. Fifteen bees were kept in each plastic jar (n_jars_=70), for a total of 1050 bees from two colonies (Supplement 3). To test for possible effect of foraging task specialization, foragers returning with pollen loads (n_P_=300) were collected from one hive and foragers returning without pollen loads (n_N_=300) from another. We presume that most bees returning without pollen loads were nectar foragers, and refer to them as such, though we did not examine their crop content (few may be water collectors or returning with empty crops). We conducted a replicate trial at a different date, in which the type of forager collected from each hive was switched (n_P_=225, and n_N_=225). The total imbibition time before release in the homing experiment was 36 h.

The second homing experiment (H2) was performed with three carbohydrate treatments at different concentrations: 8, 16, and 32% sucrose (w/w). Test bees were caught over two experimental days, again distinguishing pollen foragers from non-pollen foragers on two trial dates (Supplement 3). Twenty bees were kept per jar (n_jars_=28), with a total of 549 marked bees from two colonies. Time until release was shortened to 16 h, to avoid mortality of bees fed the low concentration of sucrose solution.

A third homing experiment (H3) was completed with four treatments: a control solution of 20% sucrose (1), or 20% sucrose enriched with 0.1% phenylalanine (2), 0.1% glycine (3), or 0.1% proline (4). With a total of 480 marked bees, fifteen individuals were kept per jar (n_jars_=32), and incubated for 20 h on top of two colonies. Each treatment was replicated eight times, including a duplication by separately testing pollen and non-pollen foragers, and the use of two colonies, others than used for H1 and H2 (Supplement 3). In H3 we tested a 20% sucrose solution, in contrast to 40% sucrose in H1. First, because amino acids taste might be masked at 40% sucrose, and second, to test within a concentration range where the carbohydrate nutrition can induce a limitation effect (see H2; survival or flight performance).

### 2.4. Data analysis

Statistics were performed with the JMP Pro software, version 10, SAS Institute Inc. Analyses were performed with general linear models, with a visual inspection of the standardized residuals to evaluate the assumptions of appropriate normality of the response variable, and normally distributed errors with homogeneous variance. In addition, parametric survival analyses (Weibull) were used to investigate flight times in the homing tests.

Forager preferences for AAs were analyzed according to the proportional consumption data by means of an Anova in which AA identity was entered as a factor with 9, 11 or 21 levels (including the controls), for nonessential, essential and all individual AAs, respectively. A post hoc Dunnett test was used to compare choice for each AA solution with the control. A potential seasonal effect on preference was also considered by testing the interaction between AAs as a factor, and the two test months March or October as a factor; this interaction was removed from the model when it was not significant. Choice among AA solutions was analyzed a second time, on AA essentiality (a 2 leveled factor; 10 essential versus 10 non-essential AAs per fence section) together with the deviation in pH value from the control (a continuous variable). The control values were excluded in this second analysis as the pH deviation was calculated relative to the control, and since the control solutions did not contain an essential or non-essential AA. Due to the double analyses on the source data, we applied a Bonferroni-correction and set the threshold for significance at α=0.025.

For the assessment of the tradeoff between AAs and carbohydrates, regressions were performed on the proportions of collected treatment solutions per fence section, considering a sucrose concentration gradient (a continuous variable of 15-24% sucrose solutions) and including the AA treatments (a 3 leveled factor: phenylalanine, glycine, control), with a post hoc Dunnett test to indicate differences between the two AAs and the control. The regression line equations allowed the extrapolation of a sucrose concentration equivalence point, which gave estimated equal consumptions of phenylalanine and glycine treatments, as compared to a control solution of 20% sucrose.

Hereafter, the phenylalanine and glycine treatment solutions, as previously extrapolated, were experimentally tested in a new artificial meadow set-up (Supplement 4), and the proportions of foraged treatment solutions per fence section were compared by means of an Anova with a Post Hoc Dunnett test, comparing the 0.1% phenylalanine and glycine solutions to the control.

For the homing experiments H1, H2 and H3, we used a factorial design, considering four factors: treatment, forager type, trial date and colony (Supplement 3). The two tested continuous response variables were the amount of test solution imbibed per bee, and the mortality during the imbibition period. The analyses were performed on mean values per replicate jar, applying post hoc Tukey tests to differentiate between levels. The return rates in homing flight tests of bees were compared by means of Chi-square tests on the numbers of returning bees versus non-returning bees, with one test for each of the factors mentioned above. The flight times in experiments were compared using the bee return data, employing survival analyses, once for every factor. P-value significance was considered after Bonferroni corrections for multiple comparisons.

## 3. Results

### 3.1. Preferences between amino acids

When tubes with 20% (w/w) sucrose solution enriched with individual AAs were offered in an artificial meadow, nectar foragers showed significant choice differences amongst the 8 nonessential AA enrichments with a control (*F*_8,63_=8.16, *P*<0.001), amongst the 10 essential AA enrichments with a control (*F*_10,77_=3.97, *P*<0.001), and also between all 20 AA enrichments with a control (*F*_20,472_=8.94, *P*<0.001) (Fig. 1A). A consistency in the order of AA choice is indicated as the consumed proportions of the 10 essential AAs, when tested alone, strongly correlated to those when all 20 AAs were tested together (regression on proportion results of AAs between tests; r^2^=0.74, *F*_1,9_=25.5, *P*<0.001). Similarly, the results on the 8 non-essential AAs alone strongly correlated to those while testing all AAs together (r^2^=0.82, *F*_1,7_=31.2, *P*<0.001). The choice among AAs was consistent between the test months March and October (interaction AA and Months: *F*_40,434_=1.22, *P*=0.24). A post hoc comparison indicated that glutamate, aspartate, histidine, cysteine and glycine were significantly disliked compared to the control (Fig. 1). Foragers preferred phenylalanine the most, although not significantly more than the control.

**Fig. 1.**
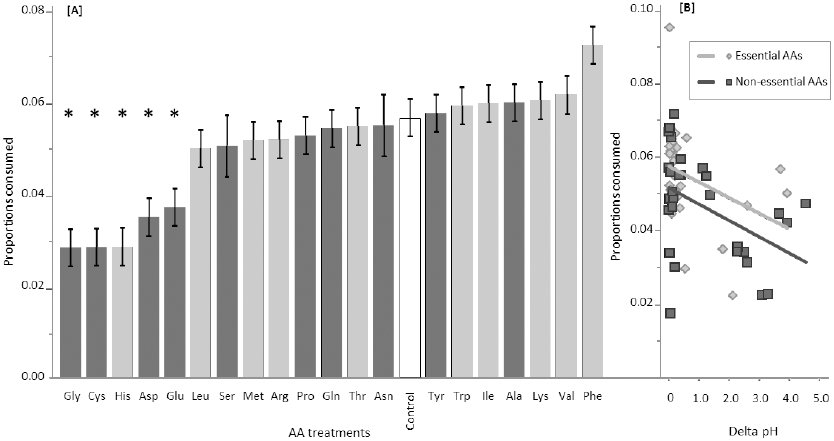
Amino acid preferences amongst free-flying foragers in an artificial meadow, according to collected proportions of AA solutions, tested simultaneously against a control. In pane A, as from the left, AAs are placed from least to most favored. The bars show mean values of collected solution amount per fence section (mean ± SE, n sections; see Supplement 1). The data represent both non-essential (dark grey) and essential (light grey) AAs. Asterisks (*) indicate AAs that were significantly different as compared to the control (white bar). Pane B shows the result of a regression between deviation in pH values (as compared to the control), and the factor essentiality. Shown are trend lines through pH deviation and mean proportional consumption data (for each AA, per trial): both pH deviation and essentiality were explanatory significant. The abbreviations for amino acids are listed in Supplement 2.

A significant preference was found for essential AA’s over non-essential AA’s (*F*_1,465_=12.17, *P*<0.001), and at the same time, the AA induced deviations in pH (absolute value relative to the control) were a significant cause for dislike (*F*_1,465_=28.54, *P*<0.001) (Fig. 1B). Overall, the acidity of the test solutions ranged between pH=2.26 (0.1% cysteine) and pH=9.33 (0.1% arginine) (Supplement 2).

### 3.2. Tradeoff between carbohydrates and amino acids

When the range of 15% to 24% sucrose solutions were offered to foragers in an artificial meadow (Fig. 2), they showed preferences for higher sucrose concentrations (*F*_1,140_= 8.05, *P*=0.005), and simultaneously phenylalanine was a significant phagostimulant, and glycine a phago-deterrent (*F*_2,140_=30.9, *P*<0.001; with posthoc significances for both glycine < control and phenylalanine > control). In Figure 2, the control regression line [y = 0.019x + 0.175] at concentration 20% indicates a consumed proportion of 0.549. At this level, the 0.1% phenylalanine regression line [y = 0.012x + 0.425] is at a concentration of 10.6% sucrose, hence at +Δ9.4% sucrose equivalence. At the same level, the 0.1% glycine regression line [y = 0.026x - 0.273] is at 31.4% sucrose, hence a - Δ11.4% sucrose equivalence. The equivalence extrapolation for phenylalanine reveals an expected AA to carbohydrate ratio of 1/94 parts w/w, and a ratio of 1/114 parts w/w for glycine.

**Fig. 2.**
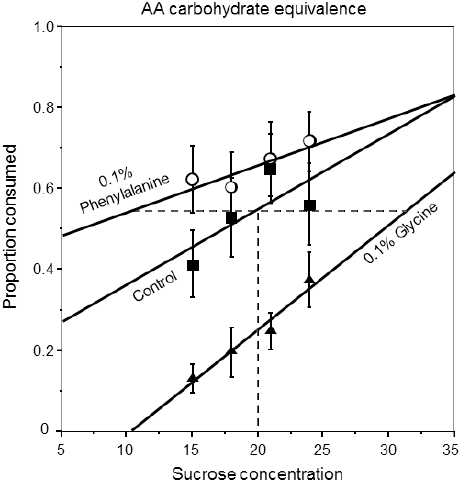
Tradeoff between AAs and carbohydrates. Bees collected sucrose solutions over a gradient of 15%, 18%, 21% and 24% sucrose, with and without AA’s. The points with error bars indicate consumed treatment solutions (mean ± SE), squares for the control, open circles for phenylalanine, and triangles for glycine. The control regression line [y = 0.019x + 0.175] at concentration 20% indicates a consumed proportion of 0.549. The horizontal dotted line indicates the extrapolated equivalence at which 0.1% AA solutions are expected to be consumed equally to the control. At this level, the 0.1% phenylalanine regression line [y = 0.012x + 0.425] is at 10.6% sucrose concentration, hence at +Δ9.4% sucrose equivalence. At the same level, the 0.1% glycine regression line [y = 0.026x - 0.273] is at 31.4% sucrose, hence at -Δ11.4% sucrose equivalence.

In two additional experiments we compensated AA presence with sucrose in order to offset the attraction and deterrence of phenylalanine and glycine, respectively (Supplement 4). In the first experiment, foragers collected dissimilar proportions of test solutions (*F*_2,51_=10.1, *P*<0.001, power=0.98 at α=0.05). An 11.6% sucrose solution with 0.1% phenylalanine was collected similar to the 20% sucrose control (*P*=0.29), whereas a 31.7% sucrose solution with 0.1% phenylalanine was collected significantly less (*P*<0.001). We therefore conducted a follow-up experiment to test sucrose compensation for the deterrent effect of glycine. Solutions were again dissimilarly collected (*F*_3,60_=25.6, *P*<0.001, power=1.00 at α=0.05), with glycine reconfirmed to be unattractive to foragers (20% sucrose + 0.1% glycine) as compared to the control (*P*=0.026). However, in this experiment the glycine effect was offset by +Δ10% sucrose (*P*=0.08), whereas glycine +Δ20% sucrose was significantly more collected than the control (*P*<0.001). Thus, in line with our extrapolations, we could confirm the AA to carbohydrate ratio (1 part phenylalanine ≈ 84 parts sucrose, and 100 parts sucrose ≤ 1 part glycine < 200 parts sucrose).

### 3.3. The effect of amino acids and carbohydrates on homing success

The results of the homing test are summarized in Tables 1, 2 and 3. The addition of 0.1% AAs in sucrose solutions did not evoke effects on treatment solution imbibition (Fig. 3), mortality and return rates (Fig. 4), or flight times (Fig. 5), either at 40% (H1) or 20% (H3) sucrose solutions. Interactions between AA treatment and other factors were not indicated.

**Fig. 3.**
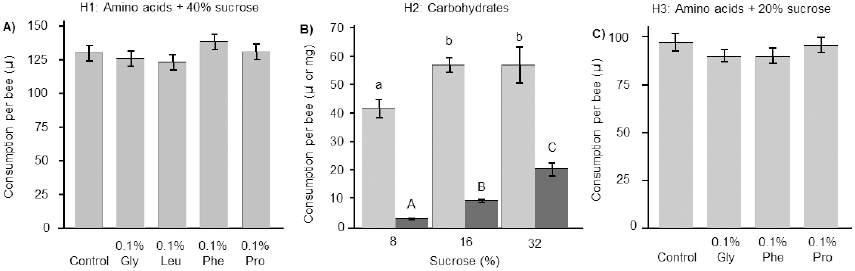
Dietary uptake (mean ± SE) of sucrose solution preceding three homing experiments, to determine whether AA and carbohydrate nutrition influence flight performance of foragers. In the first experiment (pane A: H1), forager bees were kept in jars for 36 h, whereas in the following experiments (pane B and C: H2 and H3) bees were kept in jars for 16 h and 20 h, respectively. Effect of AAs in 40% (H1) and 20% (H3) sucrose solution consumption are shown (μl bee^−1^ jar^−1^). The effect of a concentration gradient of 8%, 16%, and 32% sucrose (H2) on consumption is measured in µl solution (light bars), and given additionally as absolute amount of sucrose in mg (dark bars). Different letters represent significantly different values (P<0.05; Tukey). Notable is the indifference in imbibition of phenylalanine and glycine solutions.

**Fig 4.**
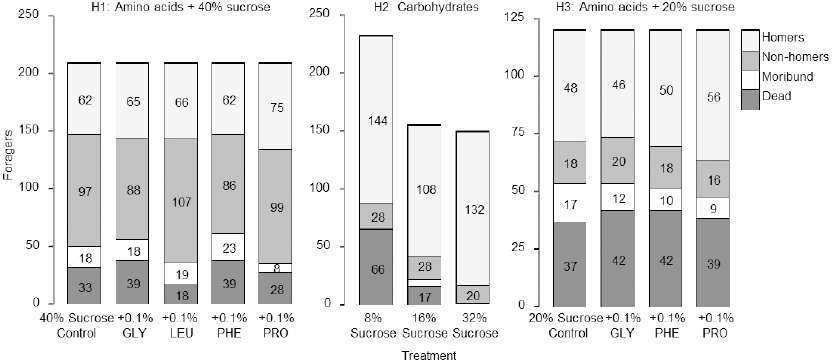
Fates of all the 2079 bees in homing experiments. Those that died during the feeding phase were classified as dead. Moribund are those that were alive at release but did not take off. Non-homers took off but did not return to the hive. Homers are those that returned to the hive. In H2 where a sucrose gradient was tested, we began with more bees in the 8% sucrose treatment, anticipating higher mortality.

**Fig 5.**
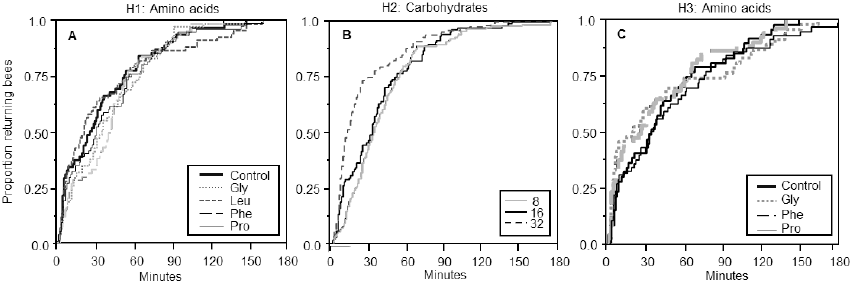
Flight time analyses of cumulative forager homecoming rates. A) Experiment H1, testing the effect of 0.1% AA nutrition in presence of 40% sucrose. B) Experiment H2, testing the effect of carbohydrate nutrition (a gradient of 8, 16 and 32% sucrose solution). C) Experiment H3, testing the effect of 0.1% AA nutrition in presence of 20% sucrose.

**Table 1:** H1 homing test results. Treatments were 40% sucrose control, and 40% sucrose enriched with 0.1% AA (glycine, leucine, phenylalanine, or proline). The imbibition test concerned 1050 bees, 70 jars, with 36 hours before testing (see Suppl. 3).

| Endpoints H1: | Uptake ($\mu\text{l bee}^{-1} \text{ jar}^{-1}$) | Mortality (mean % $\text{jar}^{-1}$) | Return rate (% bees) | Flight time (minutes) |
| --- | --- | --- | --- | --- |
| AA treatment | $F_{4,62}=1.98, P=0.11$ | $F_{4,62}=1.37, P=0.26$ | $\chi^2=1.35, Df=3, P=0.85$ | $\chi^2=5.38, Df=4, P=0.25$ |
| P & N foragers | $F_{1,62}=89.1, P<0.001^*$ | $F_{1,62}=12.6, P<0.001^*$ | $\chi^2=1.61, Df=1, P=0.20$ | $\chi^2=4.56, Df=1, P=0.033$ (ns) |
| Colony A & B | $F_{1,62}=10.4, P=0.002^*$ | $F_{1,62}=22.8, P<0.001^*$ | $\chi^2=39.4, Df=1, P<0.001^*$ | $\chi^2=0.18, Df=1, P=0.66$ |
| Trial Sep & Oct | $F_{1,62}=45.1, P<0.001^*$ | $F_{1,62}=27.5, P<0.001^*$ | $\chi^2=58.9, Df=1, P<0.001^*$ | $\chi^2=23.2, Df=1, P<0.001^*$ |
| Overall mean | 123 $\mu\text{l}$ | 157/1050 bees | 330/807 bees | median 32 min |

\* Significant differences between levels had following effect sizes: Solution uptake per forager type (P: 141 $\mu\text{l}$ > N: 107 $\mu\text{l}$ ), per colony (A: 119 $\mu\text{l}$ < B: 130 $\mu\text{l}$ ), and trial (Sept: 112 $\mu\text{l}$ < Oct: 136 $\mu\text{l}$ ). Mortality per forager type (P: 22% > N: 10%), per colony (A: 8% < B: 24%), and trial (Sept: 8% > Oct: 25%). Return rates differed per colony (A: 217 non-homers vs 224 homers; B: 206 non-homers vs 106 homers), and trial (Sept: 369 non-homers vs 169 homers; Oct: 109 non-homers vs 161 homers). Median flight time differed per trial (Sept: 18 min < Oct: 40 min), but not for forager type (P: 25 min < N: 35 min; Bonferroni corrected). No interaction between AA treatment and another factor was indicated.

**Table 2:**
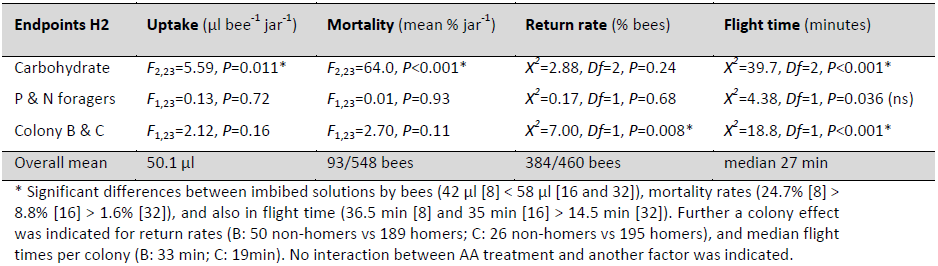
H2 homing test results. Treatments were three sucrose solution concentrations: 8, 16, and 32%. Beside pollen and nectar foragers, effects by two colonies were considered, as tested on individual trial dates. The imbibition test concerned 549 bees, 28 jars, with 16 hours before testing (see Suppl. 3).

| Endpoints H2 | Uptake ($\mu\text{l bee}^{-1} \text{ jar}^{-1}$) | Mortality (mean % $\text{jar}^{-1}$) | Return rate (% bees) | Flight time (minutes) |
| --- | --- | --- | --- | --- |
| Carbohydrate | $F_{2,23}=5.59, P=0.011^*$ | $F_{2,23}=64.0, P<0.001^*$ | $\chi^2=2.88, Df=2, P=0.24$ | $\chi^2=39.7, Df=2, P<0.001^*$ |
| P & N foragers | $F_{1,23}=0.13, P=0.72$ | $F_{1,23}=0.01, P=0.93$ | $\chi^2=0.17, Df=1, P=0.68$ | $\chi^2=4.38, Df=1, P=0.036$ (ns) |
| Colony B & C | $F_{1,23}=2.12, P=0.16$ | $F_{1,23}=2.70, P=0.11$ | $\chi^2=7.00, Df=1, P=0.008^*$ | $\chi^2=18.8, Df=1, P<0.001^*$ |
| Overall mean | 50.1 $\mu\text{l}$ | 93/548 bees | 384/460 bees | median 27 min |

\* Significant differences between imbibed solutions by bees (42 $\mu\text{l}$ [8] < 58 $\mu\text{l}$ [16 and 32]), mortality rates (24.7% [8] > 8.8% [16] > 1.6% [32]), and also in flight time (36.5 min [8] and 35 min [16] > 14.5 min [32]). Further a colony effect was indicated for return rates (B: 50 non-homers vs 189 homers; C: 26 non-homers vs 195 homers), and median flight times per colony (B: 33 min; C: 19min). No interaction between AA treatment and another factor was indicated.

**Table 3:**
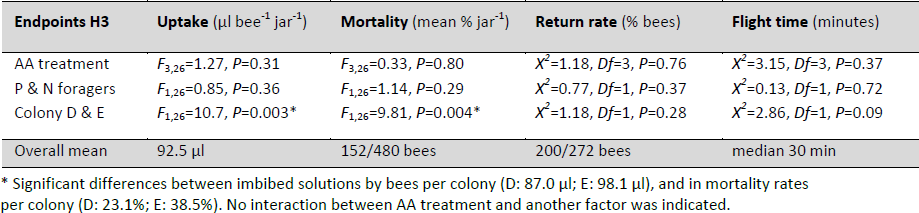
H3 homing test results. Treatments were 20% sucrose control, and 20% sucrose enriched with 0.1% AA (glycine, phenylalanine, or proline). The imbibition test concerned 480 bees, 24 jars, with 20 hours before testing. See Suppl. 3.

| Endpoints H3 | Uptake ($\mu\text{l bee}^{-1} \text{ jar}^{-1}$) | Mortality (mean % $\text{jar}^{-1}$) | Return rate (% bees) | Flight time (minutes) |
| --- | --- | --- | --- | --- |
| AA treatment | $F_{3,26}=1.27, P=0.31$ | $F_{3,26}=0.33, P=0.80$ | $\chi^2=1.18, Df=3, P=0.76$ | $\chi^2=3.15, Df=3, P=0.37$ |
| P & N foragers | $F_{1,26}=0.85, P=0.36$ | $F_{1,26}=1.14, P=0.29$ | $\chi^2=0.77, Df=1, P=0.37$ | $\chi^2=0.13, Df=1, P=0.72$ |
| Colony D & E | $F_{1,26}=10.7, P=0.003^*$ | $F_{1,26}=9.81, P=0.004^*$ | $\chi^2=1.18, Df=1, P=0.28$ | $\chi^2=2.86, Df=1, P=0.09$ |
| Overall mean | 92.5 $\mu\text{l}$ | 152/480 bees | 200/272 bees | median 30 min |

\* Significant differences between imbibed solutions by bees per colony (D: 87.0 μl; E: 98.1 μl), and in mortality rates per colony (D: 23.1%; E: 38.5%). No interaction between AA treatment and another factor was indicated.

Forager type effects were found, with nectar foragers imbibing less sucrose solutions, and having a lower mortality in H1, though this finding was not present in H2 and H3. Exclusively in H2, pollen foragers returned a notable 10 minutes earlier than nectar foragers (P: 25 min < N: 35 min; though insignificant after Bonferroni correction). Colony effects were found throughout the homing experiments, on sucrose uptake and bee mortality (between colonies A&B and D&E), return rates (between A&B and B&C), and flight times (between B&C).

The test on the effect of carbohydrate nutrition on forager homing performance (H2) showed that a relatively low volume per bee was consumed at the lowest carbohydrate concentration. A significant increase in sucrose uptake in μL and mg was observed over the sucrose gradient (*P*=0.01 and *P*<0.001, respectively, Fig. 3B). A high sucrose uptake in mg (Fig 3.) correlated with a reduction in mortality (Linear regression; *P*=0.01, *R*^2^=0.22), with 66, 17, and 1 deaths at 8%, 16% and 32% sucrose, respectively (Fig. 4-H2). The return rates of bees were not different for the three carbohydrate treatments. However, the median flight time of 14.5 minutes at the highest sucrose concentration was significantly shorter than the 35 and 36.5 minutes median it took those that received the lower sucrose concentrations (Table 2, Fig. 5B).

## 4. Discussion

As main results, we found consistent preferences of free-flying honey bee foragers for certain AAs over others, and for essential over non-essential AAs (Fig. 1). Furthermore, we found that bees were willing to pay a considerable carbohydrate premium for their AA preferences (Fig. 2). Bees were significantly attracted by 0.1% enriched phenylalanine solutions, as compared to the controls, and willing to give up 84 units sucrose for 1 unit AA. The bees were significantly deterred by glycine, but adding a 100 or more units of sucrose could resolve to offset the effect of 1 unit AA.

Roubik et al. (1995a) found the stingless bee *Melipona fuliginosa* to avoid glutamate, glycine, serine, alanine, and arginine at concentrations of 35 to 80 mM AA in 50% sucrose solutions (≈ 0.6% w/w on average), and *M. fuliginosa* valued the deterrence equal to control solutions of 20 to 40% sucrose (–Δ10% to –Δ30% sucrose). It reveals a ratio in range of 1/17 to 1/50 parts AA to carbohydrate to offset deterrence. This is less expensive than for honey bees, considering their glycine deterrence compensation with 100 or more parts sucrose.

The premiums in sucrose that bees are willing to pay for AA can be biologically significant; mortality of bees fed an 8% sucrose solution was greater than those fed 16 or 32% sucrose (Fig. 4), and it took those that survived twice as long to home back to the hive (Fig. 5B). For a honey bee colony, the caloric uptake of carbohydrates is lowered when foragers systematically prefer low sucrose nectars with phagostimulant AAs, or evade high sucrose nectars with deterrent AAs. From the plant perspective, plants can substitute expensive carbohydrates in their nectar with minute concentrations of phagostimulating AAs, or modulate pollinator visits by adding phagodeterrent AAs.

An artificial meadow assay allowed us to test the preference of free flying foragers between 20 AAs that were simultaneously present, and we found a preference for essential over non-essential AAs. Insect research has shown that an animal’s sensitivity to taste can change according to the need for nutrients due to the under-representation of nutrients evoking over-sensitivity on the receptor level (Abisgold and Simpson, 1987; Simmonds et al., 1992; Simpson et al., 1991). Perhaps, the current finding that essential AAs are more preferred than nonessentials, might relate to the yet unknown ability of forager bees to respond to specific nutritional shortcomings within their colony. We found a consistent AA preference profile between colonies at the beginning of spring growth (March) and autumn (October), but we did not specifically manipulate or assess colony needs in this study. When testing honey bees in a no-choice assay with single AAs, Inouye and Waller (1984) did not find preference effects due to essentiality of AAs. Similarly, in dual choice preference tests with *Drosophila,* Toshima and Tanimura (2012) found AA preferences unrelated to the classification of essential or non-essential, nor to other chemical properties.

The natural pH of floral nectar is reported to range between 4.2 and 8.5 (Baker, 1977). The pH of the dissolved purified AAs at 0.1% (w/w) solutions were between 2.3 and 9.3, which lies beyond the natural range in nectar (Supplement 2). We found that 0.1% AA could affect nectar pH, and that foragers were sensitive to the pH (Fig. 1B). The statistical model showed independent effects on bee preferences of whether AAs were essential or not and of their pH. Furthermore, the pH of glycine and phenylalanine enriched solutions were similar, yet the former was consistently disliked by the bees whereas the latter was consistently preferred (Fig. 2). Thus, pH affects forager choice but only partially explains preferences between AAs. The pH of the resulting solution should always be considered in taste or preference studies that dissolve purified AAs in solution.

In our study, bees could use both olfaction and gustation to discriminate between AA solutions. Linander et al. (2012) showed that bees could discriminate between the odor of single dissolved AAs and the control solvent, specifically aspartate, cysteine, proline, tryptophan, and tyrosine. A conversion by molecular weight indicates our AA concentrations in range of 5.3 – 14.3 mM (supplement 2). This is one order of magnitude smaller than the test concentration of 100 mM as used by Linander et al. (2012). Nonetheless, the discrimination between cysteine and aspartate from the control in our experiments (Fig. 1) could thus have been facilitated by olfactory cues. We found that free-flying foragers could also discriminate phenylalanine and glycine from the control (Fig. 2). This discrimination may have relied on gustation. Although bees are able to discriminate proline, tryptophan and tyrosine from the control by scent at 100mM (Linander et al., 2012), we did not find preference effects for these AAs at the tested concentration of 0.1% (<10mM).

Honey bees show a significant phagostimulatory effect of phenylalanine and a phagodeterrent effect of glycine (Fig. 2), which is consistent with the findings of Inouye and Waller (1984). However, considering the results in Figure 1, the choice among AAs for which bees did not show extreme preferences was not always consistent with other studies. For example, proline at concentrations similar to those that we studied, has generally been reported to be attractive to bees (Bertazzini et al., 2010; Carter et al., 2006; Inouye and Waller, 1984). Similarly, bees in our study favored alanine, but not significantly more than the control; Bertazzini et al. (2010) and Inouye and Waller (1984) found alanine to be significantly preferred to the control. Different preferences of bees between studies may be due to methodology (e.g., choice vs no-choice, free-flying vs caged), or differences in sensitivity thresholds of different honey bee strains. Such inconsistencies are more likely to show for AAs that evoke less pronounced preferences; AAs at the extreme ends seem to be valued more consistently.

Of course, preference and deterrence effects are also dependent on AA concentrations. However, AAs in nectar vary greatly within plants and between species within several orders of magnitude (Baker, 1977; Nepi et al., 2012; Weiner et al., 2010). Therefore, we chose to standardize our experimental setup, and test all AAs at the same concentration of 0.1% (w/w), which lies within the higher boundary of the natural range. It is for instance comparable to *Lantana camara* nectar, with 16 mM total AA (an approximate 0.1% w/w); a plant visited by bees and many butterflies and described to have a relative high amino acid concentration (Alm et al., 1990).

In addition to honey bees, AA preferences are reported for tropical stingless bees (Gardener et al., 2003; Roubik et al., 1995b), solitary bees (Weiner et al., 2010), butterflies (Erhardt and Rusterholz, 1998), ants (Bluthgen and Fiedler, 2004a, b; Wada et al., 2001), flies (Potter and Bertin, 1988), and nectivorous bats (Rodriguez-Pena et al., 2013). Differences in taste perception between diverse pollinators likely affect nectar perception and may be a factor in how plants bias pollinator visits, thereby affecting gene flow within and amongst plant populations (Nepi et al., 2012).

AA concentrations in nectar are low relative to pollen, thereby suggesting a limited role in bee nutrition. For pollen, it has been described that plants visited by oligolectic insects have a lower pollen quality, for instance in the ratio of essential to non-essential AAs, as compared to plants visited only by non-oligolectic insects (Weiner et al., 2010). The noted pollen quality differentiation was however not reflected in the free amino acids content of the pollens. Thus, since free amino acids are the most likely cue for a potential taste-based evaluation of AA nutrition, it is not likely that pollinators respond to nutritional quality based on AA taste alone. Also, most of the 20 AAs tested in the current study evoked neutral to mild deterrent responses in honey bees, thus it seems unlikely that bees respond to AAs in nectar as a cue to assess nutritional quality.

A substantial question within the current study addressed if AAs in nectar may affect the physiology and performance of bees. As mentioned, proline may be directly used as a minor source of flight fuel (Barker and Lehner, 1972; Carter et al., 2006; Micheu et al., 2000; Mollaei et al., 2013). However, the study by Mollaei et al. (2013) showed no notable honey bee flight muscle stimulation at 0.1% proline treatment. In our homing studies proline at 0.1% did also not evoke an effect, neither on the return rate, nor the median flight time of bees (Table 1 and 3).

In contrast to the consistent preferences exhibited by free-flying bees, caged foragers in a no-choice context imbibed equally from sucrose solution or sucrose solution enriched with AAs (Fig. 3). An dietary enrichment with either Phenylalanine, Glycine, Leucine or Proline did not affect their homing performance (Fig. 5A) or other physiological measures of performance (Table 1 and 3). However, physiology and performance were directly influenced by carbohydrate nutrition (H2): foragers fed with lower sucrose concentrations took longer to come back to the hive (Fig. 4). This can have important methodological implications for homing studies that evaluate the effect of substances that may affect the imbibition of sugar solution.

Forager task specialization affects various physiological and behavioral traits (Nelson et al., 2007; Page, 2013; Robinson and Vargo, 1997). In one of our homing tests (H1) pollen foragers imbibed more test solution than non-pollen foragers (Table 1). Homing ability is an effective assay in eco-toxicological tests, and the amount of test solution imbibed would affect exposure to toxicants. Our study suggests that controlling for task specialization could reduce experimental variability.

We have presented a novel and useful bioassay by means of an artificial meadow set-up. We illustrate high test sensitivity, in which steps of only Δ3% sucrose concentration show a clear gradient effect, in addition to detecting an AA treatment effect (Fig. 2). This simple setup could be useful for tackling additional questions about bee perception of nectar components. Specifically for AAs, we tested them singly, but Alm et al. (1990) found that honey bees consumed more and longer from nectar sources which were enriched with a mixture of AAs, compared to nectar without AAs. This supports the hypothesis that the AAs in nectar contribute to pollinator attraction. Higher stimulatory effects of mixtures of AAs over single AAs have been frequently reported also for ants (Lanza, 1988; Lanza and Krauss, 1984). The artificial meadow set-up would be useful for further research into the complexity of AA attraction of mixtures compared to single AAs.

In conclusion, we found three variables that explain observed AA preferences of honey bee nectar foragers: AA identity, pH, and essentiality. We identified sucrose equivalents that foragers are willing to exchange for phagostimulant or phagodeterrent AAs. Flight performance of foraging bees was affected by nutritional state: foragers were directly affected by carbohydrate levels, but not directly by either proline, leucine, phenylalanine or glycine, nor indirectly by their excitatory or deterrent tastes. Both pH and AAs were detected by the bees in artificial nectar at natural concentrations. Thus, such preferences by pollinators could affect floral evolution, shaping the biodiversity in natural ecosystems, and could affect pollination services in agricultural landscapes.

## Acknowledgements

This research was funded jointly by a grant from BBSRC, Defra, NERC, the Scottish Government and the Wellcome Trust, under the Insect Pollinators Initiative. We thank Tania Masci, Tal Erez and Yael Katz for assisting in marking bees and recording data, and for Haim Kalev for apiary management. We also thank Claudio Rolli, Marleen de Blécourt, two anonymous reviewers and the editor for their valuable and constructive comments on the MS.

## Supplement 1

**S1:**
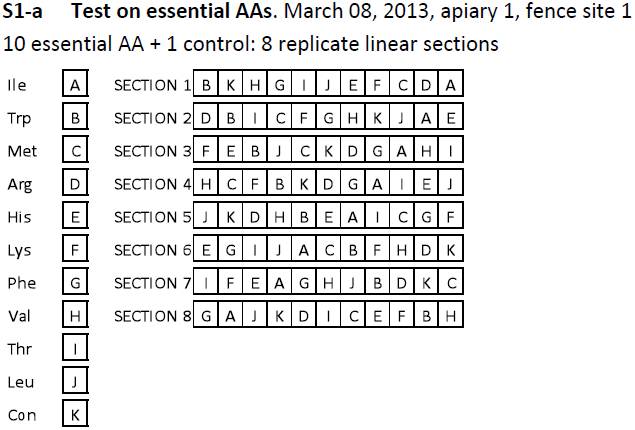
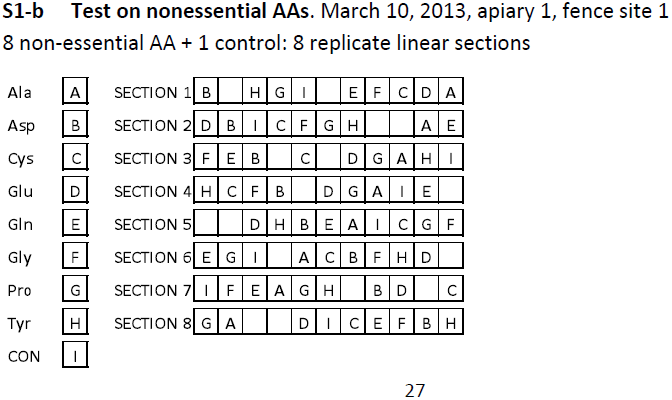
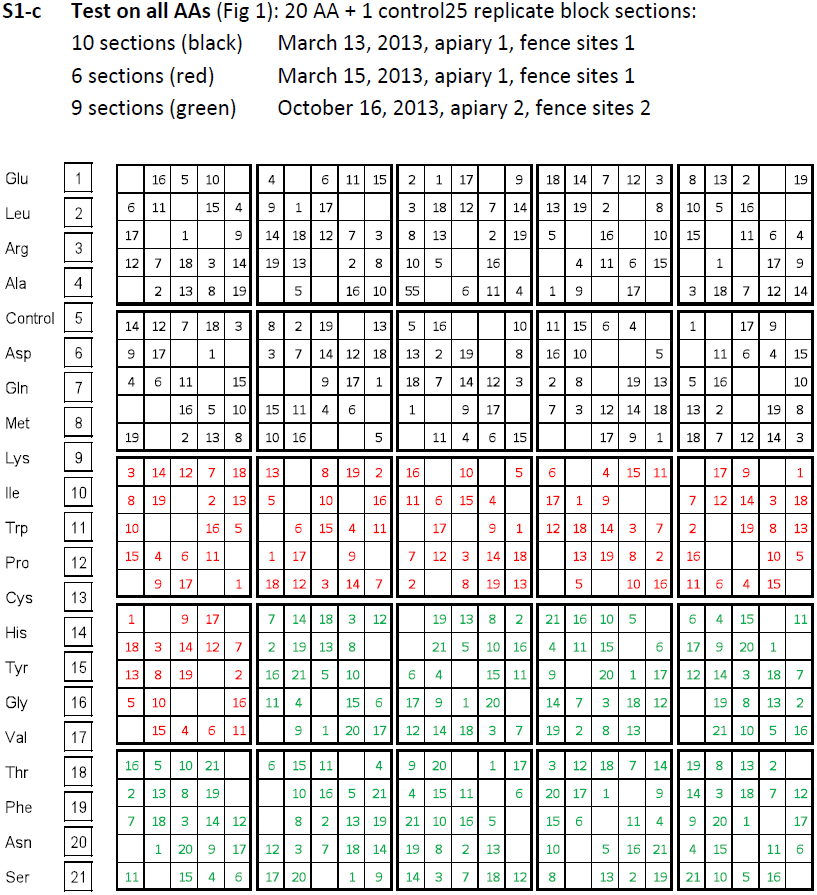
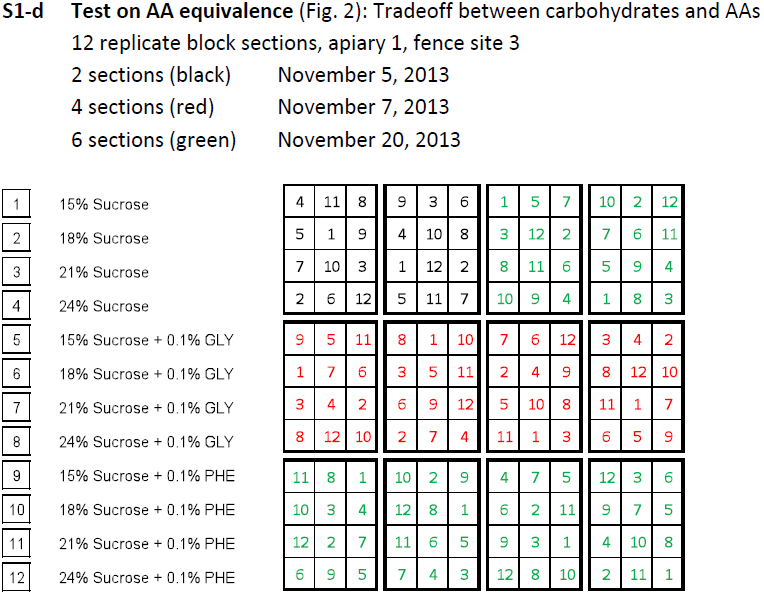
EXPERIMENTAL DESIGNS OF FENCE TESTS: AA PREFERENCE AND EQUIVALENCE The experimental designs of the preference tests of AA on free flying foragers. Treatment solutions were offered to the bees at fence sites close to apiaries. The design follows a Latin square considering that a treatment position in a replicate section is represented only once, so that potential spatial effects are equally balanced within the experimental setups. Three geographically different fence sites were used, on two of our apiaries. Each apiary contained 20-30 hives which were exchanged from time to time. The tests were performed over several months.

## Supplement 2

**S2:**
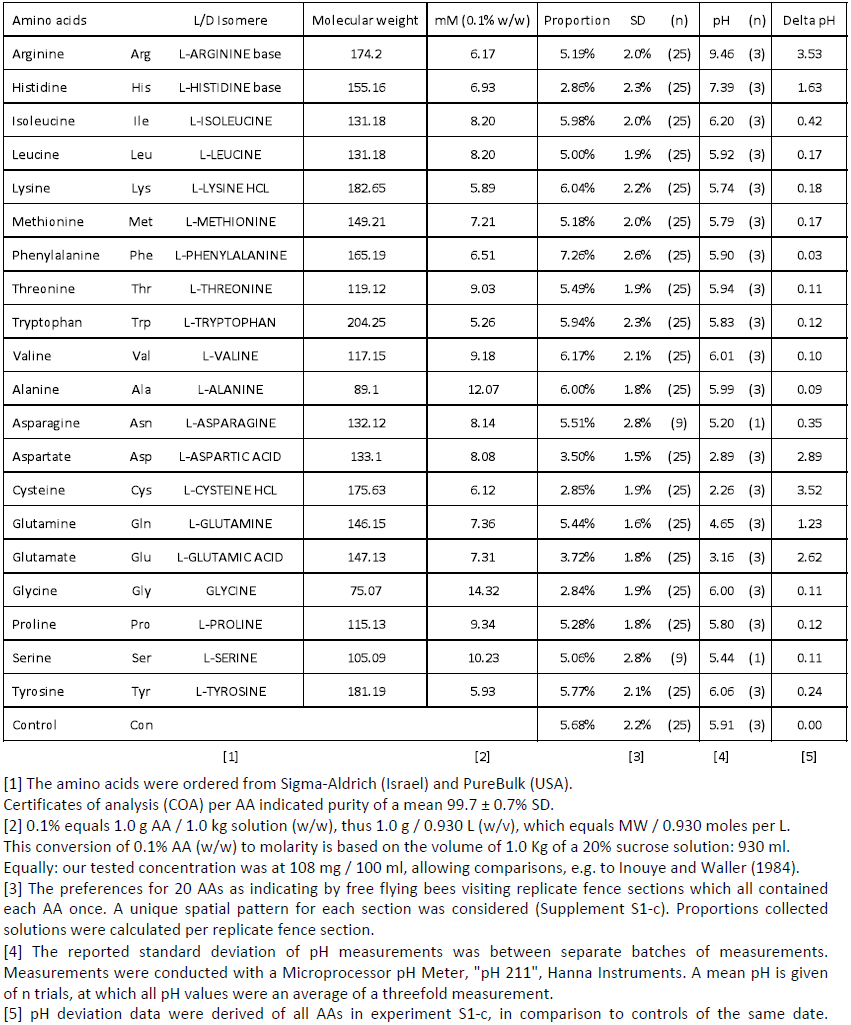
TABLE AMINO ACID INFORMATION (Fig. 1; Supplement S1-c) Background information on amino acids for tests on preferences by nectar foragers

| Amino acids | L/D Isomere |  | Molecular weight | mM (0.1% w/w) | Proportion | SD | (n) | pH | (n) | Delta pH |
| --- | --- | --- | --- | --- | --- | --- | --- | --- | --- | --- |
| Arginine | Arg | L-ARGININE base | 174.2 | 6.17 | 5.19% | 2.0% | (25) | 9.46 | (3) | 3.53 |
| Histidine | His | L-HISTIDINE base | 155.16 | 6.93 | 2.86% | 2.3% | (25) | 7.39 | (3) | 1.63 |
| Isoleucine | Ile | L-ISOLEUCINE | 131.18 | 8.20 | 5.98% | 2.0% | (25) | 6.20 | (3) | 0.42 |
| Leucine | Leu | L-LEUCINE | 131.18 | 8.20 | 5.00% | 1.9% | (25) | 5.92 | (3) | 0.17 |
| Lysine | Lys | L-LYSINE HCL | 182.65 | 5.89 | 6.04% | 2.2% | (25) | 5.74 | (3) | 0.18 |
| Methionine | Met | L-METHIONINE | 149.21 | 7.21 | 5.18% | 2.0% | (25) | 5.79 | (3) | 0.17 |
| Phenylalanine | Phe | L-PHENYLALANINE | 165.19 | 6.51 | 7.26% | 2.6% | (25) | 5.90 | (3) | 0.03 |
| Threonine | Thr | L-THREONINE | 119.12 | 9.03 | 5.49% | 1.9% | (25) | 5.94 | (3) | 0.11 |
| Tryptophan | Trp | L-TRYPTOPHAN | 204.25 | 5.26 | 5.94% | 2.3% | (25) | 5.83 | (3) | 0.12 |
| Valine | Val | L-VALINE | 117.15 | 9.18 | 6.17% | 2.1% | (25) | 6.01 | (3) | 0.10 |
| Alanine | Ala | L-ALANINE | 89.1 | 12.07 | 6.00% | 1.8% | (25) | 5.99 | (3) | 0.09 |
| Asparagine | Asn | L-ASPARAGINE | 132.12 | 8.14 | 5.51% | 2.8% | (9) | 5.20 | (1) | 0.35 |
| Aspartate | Asp | L-ASPARTIC ACID | 133.1 | 8.08 | 3.50% | 1.5% | (25) | 2.89 | (3) | 2.89 |
| Cysteine | Cys | L-CYSTEINE HCL | 175.63 | 6.12 | 2.85% | 1.9% | (25) | 2.26 | (3) | 3.52 |
| Glutamine | Gln | L-GLUTAMINE | 146.15 | 7.36 | 5.44% | 1.6% | (25) | 4.65 | (3) | 1.23 |
| Glutamate | Glu | L-GLUTAMIC ACID | 147.13 | 7.31 | 3.72% | 1.8% | (25) | 3.16 | (3) | 2.62 |
| Glycine | Gly | GLYCINE | 75.07 | 14.32 | 2.84% | 1.9% | (25) | 6.00 | (3) | 0.11 |
| Proline | Pro | L-PROLINE | 115.13 | 9.34 | 5.28% | 1.8% | (25) | 5.80 | (3) | 0.12 |
| Serine | Ser | L-SERINE | 105.09 | 10.23 | 5.06% | 2.8% | (9) | 5.44 | (1) | 0.11 |
| Tyrosine | Tyr | L-TYROSINE | 181.19 | 5.93 | 5.77% | 2.1% | (25) | 6.06 | (3) | 0.24 |
| Control | Con |  |  |  | 5.68% | 2.2% | (25) | 5.91 | (3) | 0.00 |

[1] The amino acids were ordered from Sigma-Aldrich (Israel) and PureBulk (USA).
Certificates of analysis (COA) per AA indicated purity of a mean $99.7 \pm 0.7\%$ SD.
[2] 0.1% equals 1.0 g AA / 1.0 kg solution (w/w), thus 1.0 g / 0.930 L (w/v), which equals MW / 0.930 moles per L.
This conversion of 0.1% AA (w/w) to molarity is based on the volume of 1.0 Kg of a 20% sucrose solution: 930 ml.
Equally: our tested concentration was at 108 mg / 100 ml, allowing comparisons, e.g. to Inouye and Waller (1984).
[3] The preferences for 20 AAs as indicating by free flying bees visiting replicate fence sections which all contained each AA once. A unique spatial pattern for each section was considered (Supplement S1-c). Proportions collected solutions were calculated per replicate fence section.
[4] The reported standard deviation of pH measurements was between separate batches of measurements. Measurements were conducted with a Microprocessor pH Meter, "pH 211", Hanna Instruments. A mean pH is given of n trials, at which all pH values were an average of a threefold measurement.
[5] pH deviation data were derived of all AAs in experiment S1-c, in comparison to controls of the same date.

## Supplement 3

**S3:**
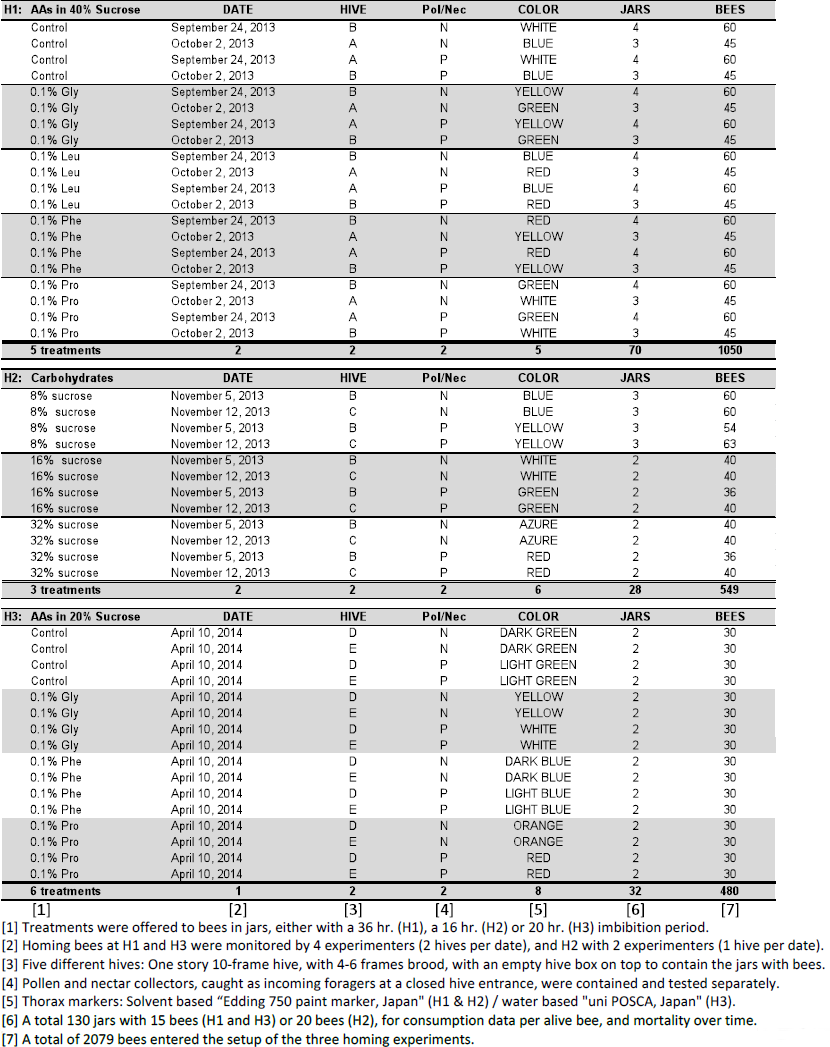
TABLE AMINO ACID INFORMATION (Fig. 1; Supplement S1-c) Experimental designs of three homing experiments, as described in the materials and methods section. The tests aimed to monitor AA nutrition effects on forager flight (H1 and H3) considering AA-induced alterations in carbohydrate uptake. Separately, carbohydrate uptake effects were tested in the second experiment (H2).

## Supplement 4

**S4:**
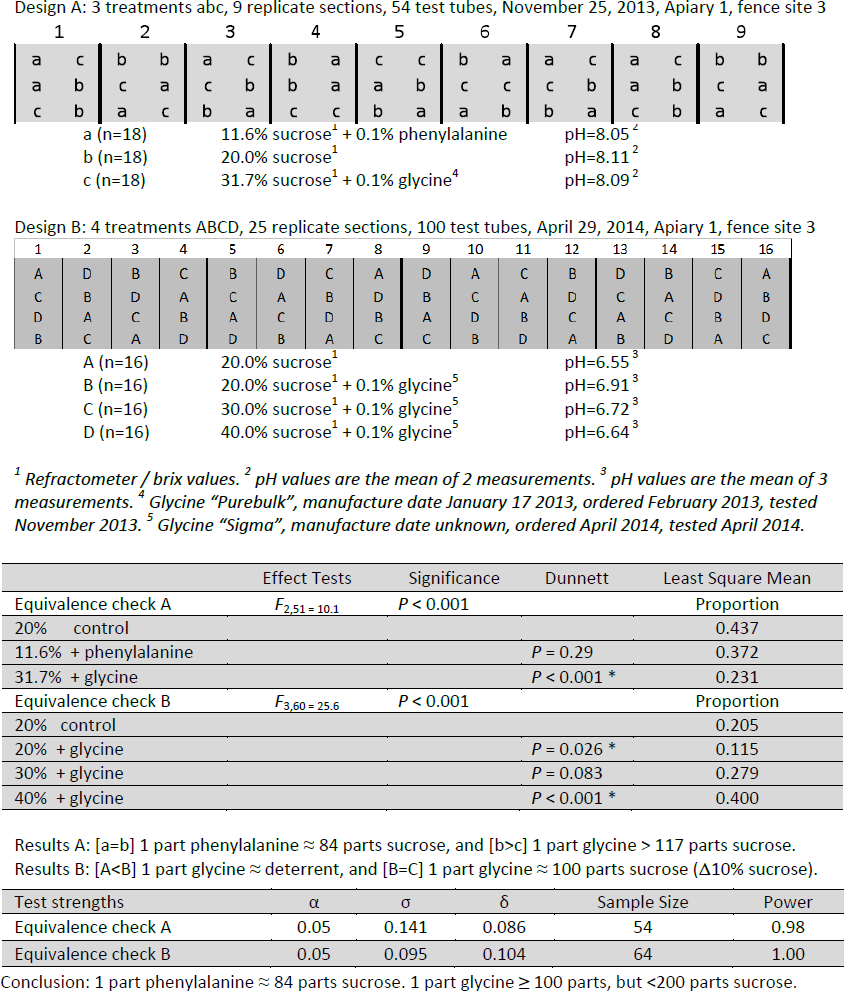
EQUIVALENCE EXPERIMENTS For the AAs phenylalanine and glycine, an extrapolation assessed the equivalence of 0.1% AA to % sucrose (S1-d). The extrapolated points were beyond the tested gradient, thus additional experiments were performed to verify the expected equivalence: 20% sucrose ≈ 10.6% sucrose + 0.1 % phenylalanine ≈ 31.4% sucrose + 0.1% glycine

